# Face identity selectivity in the deep neural network and human brain

**DOI:** 10.1101/2021.05.12.443829

**Authors:** Jinge Wang, Runnan Cao, Nicholas J Brandmeir, Xin Li, Shuo Wang

## Abstract

A central challenge in face perception research is to understand how neurons encode various face identities. However, this challenge has not been met largely due to the lack of simultaneous access to the activity of the entire face processing neural network as well as the lack of a comprehensive multifaceted model that is able to characterize a large number of facial features. In this study, we address this challenge by conducting *in silico* experiments using a deep neural network (DNN) capable of face recognition with a diverse array of stimuli. We identified a subset of DNN neurons selective to face identities, and these identity-selective neurons demonstrated generalized discriminability to novel faces not involved in the training and in many different styles. Visualization of the network explained the response of the DNN neurons and manipulation of the network confirmed the importance of identity-selective neurons in face recognition. Importantly, using our human single-neuron recordings, we directly compared the response of artificial neurons with 490 real human neurons to the same stimuli and found that artificial neurons did share a similar representation of facial features as human neurons. We also observed a novel region-based feature coding mechanism in DNN neurons as in human neurons, which may explain how the DNN performs face recognition. Together, by directly linking between artificial and human neurons, our results shed light on how human neurons encode face identities.

## Introduction

The ability to identify and recognize faces is one of the most important cognitive functions in social communications. Primates have a dedicated neural system to process faces. Neurons that are selectively responsive to faces (i.e., face-selective neurons) have been observed in a distributed network of brain areas, notably including the face patches in the temporal cortex [1] (shown in monkeys) as well as the amygdala and hippocampus [2, 3] (shown in both monkeys and humans). In particular, there are two extreme hypotheses about how neurons encode and represent faces. The exemplar-based model posits that faces are represented by highly selective, sparse, but visually invariant neurons [4–7]. This model has been supposed by single-neuron recordings in the human amygdala, hippocampus, and other parts of the medial temporal lobe (MTL). The feature-based model posits that faces are represented by simultaneous activation of a broad and distributed population of neurons and each neuron responds to many pictures with similar basic features [8, 9]. This model has been supported by single-neuron recordings in the monkey inferotemporal (IT) cortex [10–13]. Although exemplar-based and feature-based models are not mutually exclusive because both types of neurons have been observed in different brain regions, it remains unclear how to bridge these prior findings and how the brain transitions from one model to the other, which is largely due to the lack of simultaneous access to the activity of the entire face processing neural network (i.e., the whole population of neurons from all brain regions involved in face processing) [14].

To address this limitation, in this study, we exploit the opportunity to analyze the activity of all neurons from an artificial neural network (ANN) dedicated to face recognition. Importantly, we were able to compare the results directly with recordings from the human neurons, which will shed light on how the human brain encode face identities. Rapid advances in deep neural networks (DNN) have offered new opportunities for studying face perception by providing computational proxy models. State-of-the-art DNNs such as the VGG-face [15] and DeepFace [16] have achieved excellent face recognition performance and even outperformed humans. These DNNs are biologically inspired and therefore have the potential to successfully provide insight into the underlying mechanisms of brain functions, especially with respect to the perception and recognition of visual stimuli such as faces. Existing work at the intersection of DNNs and face perception can be broadly classified into two categories: *face reconstruction* (decoding models) and *face recognition* (encoding models). The former includes the reconstruction of faces from fMRI patterns [17, 18] (see [19, 20] for more general natural image reconstruction). The latter includes a flurry of literature on the convergent evolution of face spaces across DNN layers and human face-selective brain areas [21], the neurally plausible efficient inverse graphics model for face processing [22], and spontaneous generation of face recognition in untrained DNNs [23].

The present study continues from our recent work showing feature-based encoding of face identities in the human MTL using DNN-extracted visual features [24] (an encoding-related study). The motivation for the present study is largely two-fold. First, the sparse coding hypothesis for face recognition has been inspected recently under the framework of DNN. It has been shown that identity, gender, and viewpoint information all contributes to individual unit responses [25, 26], similar to the neuronal coding of facial attributes in the primate brain [9, 24, 27]. Experimentally, it is difficult to comprehensively characterize the tuning properties of neurons from the human brain to a large number of facial attributes; therefore, in this study, we will conduct *in silico* experiments to probe the tuning properties of DNN neurons using a diverse array of stimuli, which will in turn shed light on the tuning properties of human neurons. Second, several lines of research in DNN (e.g., dropout [28], neural architecture search [29]) have shown supporting evidence about the so-called *lottery ticket hypothesis* [30] (i.e., among all possible fully-connected and feedforward architectures, the winning ticket is a sparse subnetwork). This hypothesis has important biological implications into both energy efficiency and generalization properties, but there is still a missing parallel and convergent connection between the sparse coding hypothesis in neuroscience [31] and the lottery ticket hypothesis in deep learning [30]. To fill in this gap, in this study we used a DNN as a proxy model for studying the tuning properties of neurons that encode face identities. For the first time, our present study will directly compare and link the sparse coding of face identities between human and DNN neurons.

## Results

### Identity-selective neurons

We used 500 natural face images of 50 celebrities (10 faces per identity) to elicit response from a pre-trained deep neural network (DNN) VGG-16 trained to recognize faces (see **Fig. 1A** for DNN architecture). The pre-trained DNN had an accuracy of 94.2%±2.3% (mean±SD across identities) in identity recognition after fine-tuning. We identified a subset of DNN neurons that showed a significantly unequal response to different identities (one-way ANOVA of activation for each DNN neuron: P < 0.01; **Fig. S1A**), and we refer to this population of neurons as identity-selective neurons (**Fig. 1B**). The percentage of identity-selective neurons was higher in the later DNN layers (**Fig. 1B**) because the later DNN layers were closer to the output (note that the output was the identity of the input face). Interestingly, we observed that a substantial amount of neurons were selective to multiple identities (referred to here as multiple-identity [MI] neurons; **Fig. 1C**; see **Methods** and **Fig. S1A** for selection procedure), consistent with prior studies with direct recordings from single neurons in the human brain [24, 32]. Compared to neurons that were selective to a single identity (referred to here as single-identity [SI] neurons), the percentage of MI neurons increased in the later DNN layers (**Fig. 1C**). Furthermore, some identities were encoded by more SI and MI neurons than the other identities (**Fig. S1B**).

**Fig. 1.**
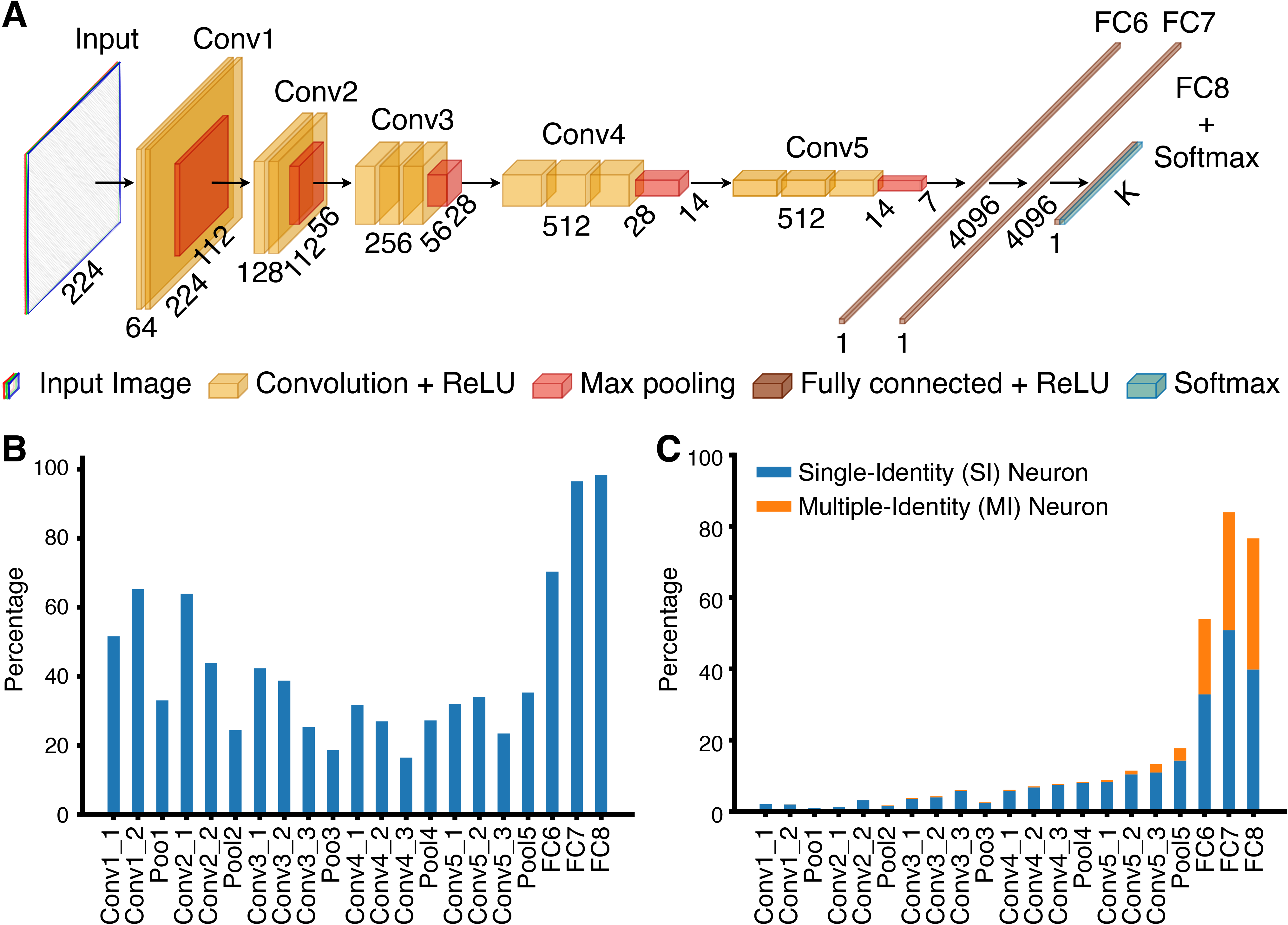
Identity-selective neurons in a pre-trained VGG-16 deep neural network (DNN). **(A)** Structure of the VGG-16 DNN. The convolutional neural network (CNN) consisted of a feature extraction section (13 convolutional layers) and a classification section (3 fully connected [FC] layers). The feature extraction section was consistent with the typical architecture of a CNN. A 3×3 filter with 1-pixel padding and 1-pixel stride was applied to each convolutional layer, which followed by Rectified Linear Unit (ReLU) operation. Every convolutional block was followed by a max-pool operation with a stride of 2. There were 3 FC layers in each classification section: the first two had 4096 channels each, and the third performed an *K*-way classification. Each FC layer was followed by a ReLU and 50% dropout to avoid overfitting. A nonlinear Softmax operation was applied to the final output of VGG-16 network to make the classification prediction of 50 identities. **(B)** Percentage of identity-selective neurons for each DNN layer. **(C)** Percentage of single-identity (SI; blue) and multiple-identity (MI; orange) neurons for each DNN layer.

### Identity-selective neurons demonstrated generalized selectivity to face identities

We next analyzed the selectivity properties of identity-selective neurons. We used a support vector machine (SVM) to assess to what extent neurons from a specific DNN layer could distinguish the input stimuli. First, with the original stimuli used to select identity-selective neurons, we found that the discriminability of all DNN neurons for face identities increased in the later DNN layers (**Fig. 2A**). This was expected because the later DNN layers were closer to the output and contained more information about face identities. Notably, such discriminability was primarily driven by identity-selective neurons and non-identity-selective neurons alone could not discriminate different identities (**Fig. 2A**). Interestingly, identity-selective neurons had even better discriminability than all neurons in the earlier DNN layers (**Fig. 2A**).

**Fig. 2.**
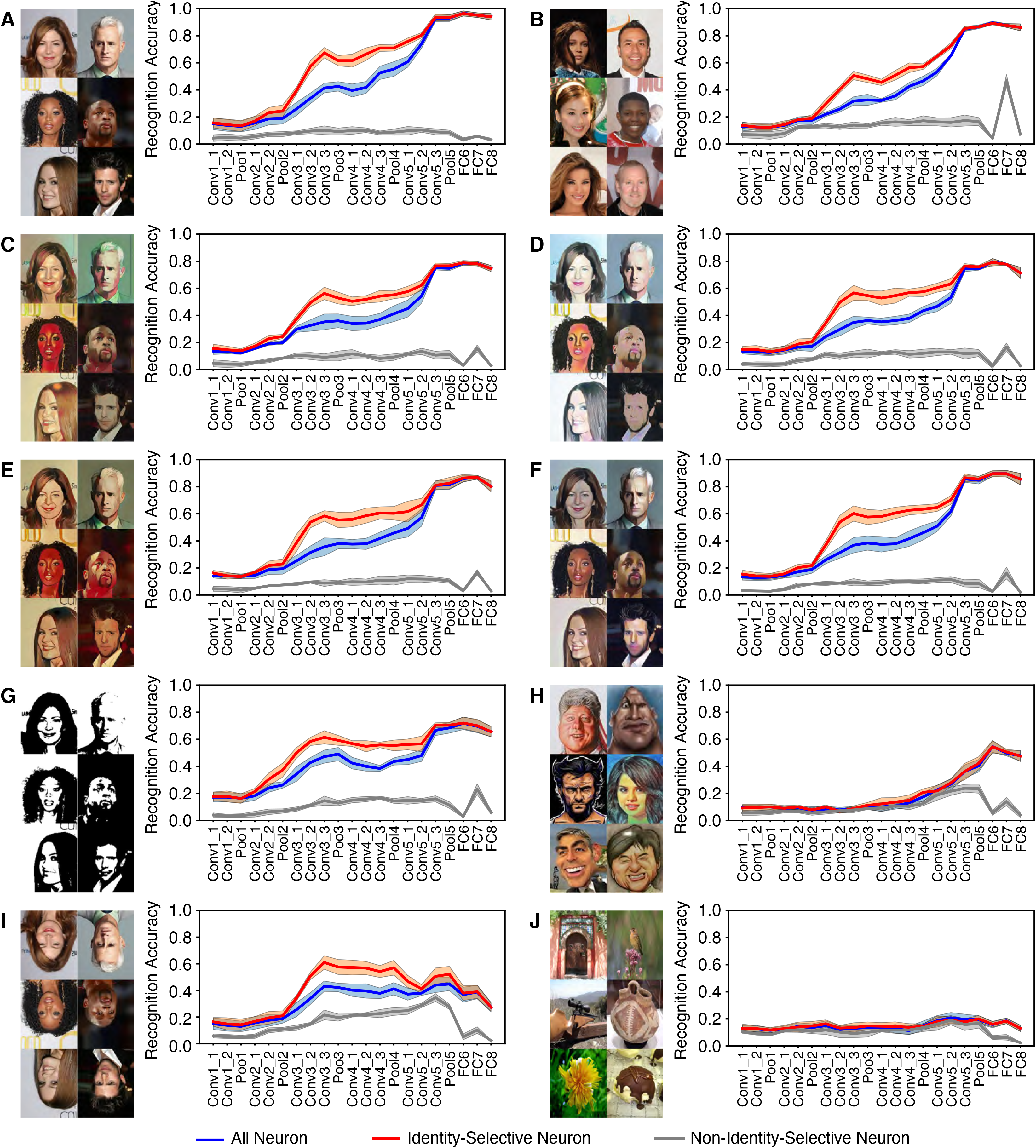
Selectivity properties of identity-selective neurons. **(A)** Original faces used to identify identity-selective neurons. **(B)** Faces from a different set of 50 identities randomly selected from the CelebA database. **(C)** Original faces in the cartoon style Hayao. **(D)** Original faces in the cartoon style Hosoda. **(E)** Original faces in the cartoon style Paprika. **(F)** Original faces in the cartoon style Shinkai. **(G)** Original faces in the Mooney style. **(H)** A different set of celebrity caricature faces. **(I)** Original faces in inversion. **(J)** A set of non-face objects selected from the ImageNet stimuli. Identity recognition accuracy is shown for each deep neural network (DNN) layer. Error shade denotes one standard deviation across 5-fold cross validation. Blue: all neurons from each DNN layer. Red: identity-selective neurons. Gray: non-identity-selective neurons.

Second, we found that the DNN could discriminate identities in a different set of celebrity faces (**Fig. 2B**) as well as the original celebrity faces transformed to various cartoon styles (**Fig. 2C-F**), and the response profile was similar to that with the original stimuli (although cartoon faces had overall reduced discrimination accuracy): identity-selective neurons primarily drove the discrimination and even had better discriminability than all neurons in the earlier DNN layers whereas non-identity-selective neurons could not discriminate identities in all these tests.

Third, we found that identity-selective neurons could still discriminate face identities even with very limited information in faces (**Fig. 2G**). Although the accuracy was reduced to discriminate these low-information two-tone Mooney faces (generated from the original celebrity faces), a similar pattern of response was observed with identity-selective neurons primarily driving the discrimination.

Fourth, we found that identity-selective neurons could discriminate a different set of celebrity caricature faces in an exaggerated cartoon style (**Fig. 2H**). Although the overall accuracy was reduced, identity-selective neurons still played the dominant role in discriminating the identities and non-identity-selective neurons did not contribute to the discrimination. It is also worth noting that identity-selective neurons as well as all neurons could not discriminate face identities any more in earlier layers (**Fig. 2H**).

Fifth, we found the DNN could still discriminate inverted faces (**Fig. 2I**), although the accuracy was reduced, consistent with impaired discrimination of inverted face in humans [33]. Such discrimination was again driven by identity-selective neurons. However, we found that the DNN (all neurons, identity-selective neurons, and non-identity-selective neurons) could barely discriminate non-face object categories (**Fig. 2J**). Therefore, the response could only be generalized within faces.

Lastly, we found that DNN neurons selective to identities from one race (e.g., Caucasian) could also discriminate face identities from other races, suggesting that the DNN and identity-selective neurons had cross-race generalizability. Furthermore, we found that DNN neurons selective to identities from one gender (e.g., male) could also discriminate face identities from the other gender, suggesting that the DNN and identity-selective neurons had cross-gender generalizability. In addition, we found that combined SI and MI neurons (i.e., a subset of identity-selective neurons selected by an additional criterion that the response for certain identities stood out from the global mean response; see **Methods** and **Fig. S1A**) demonstrated even stronger discriminability of face identities (**Fig. S1C**).

Together, our results showed that identity-selective neurons played a general and critical role in discriminating face identities under various circumstances, whereas non-identity-selective neurons could discriminate face identities in none of the circumstances. Therefore, our results suggested that although the discriminability varied as a function of the level of information contained in the stimuli, a subset of DNN neurons were consistently involved in the face identity discrimination and these neurons demonstrated generalized selectivity to face identities.

### DNN visualization explained the role of identity-selective neurons in face recognition

We next visualized the response of identity-selective vs. non-identity-selective neurons in order to understand why identity-selective neurons but not non-identity-selective neurons played a general and critical role in face recognition. First, we found that identity-selective neurons indeed corresponded to the critical visual features of the stimuli (**Fig. 3A**). Second, when we constructed a two-dimensional stimulus feature space using t-distributed stochastic neighbor embedding (t-SNE) feature reduction for each DNN layer, we found that face identities clustered in the feature space constructed by identity-selective neurons but not non-identity-selective neurons (**Fig. 3B**), confirming that identity-selective neurons could discriminate face identities. Together, DNN visualization revealed that identity-selective neurons encoded critical stimulus features so that they embodied a general discriminability of face identities.

**Fig. 3.**
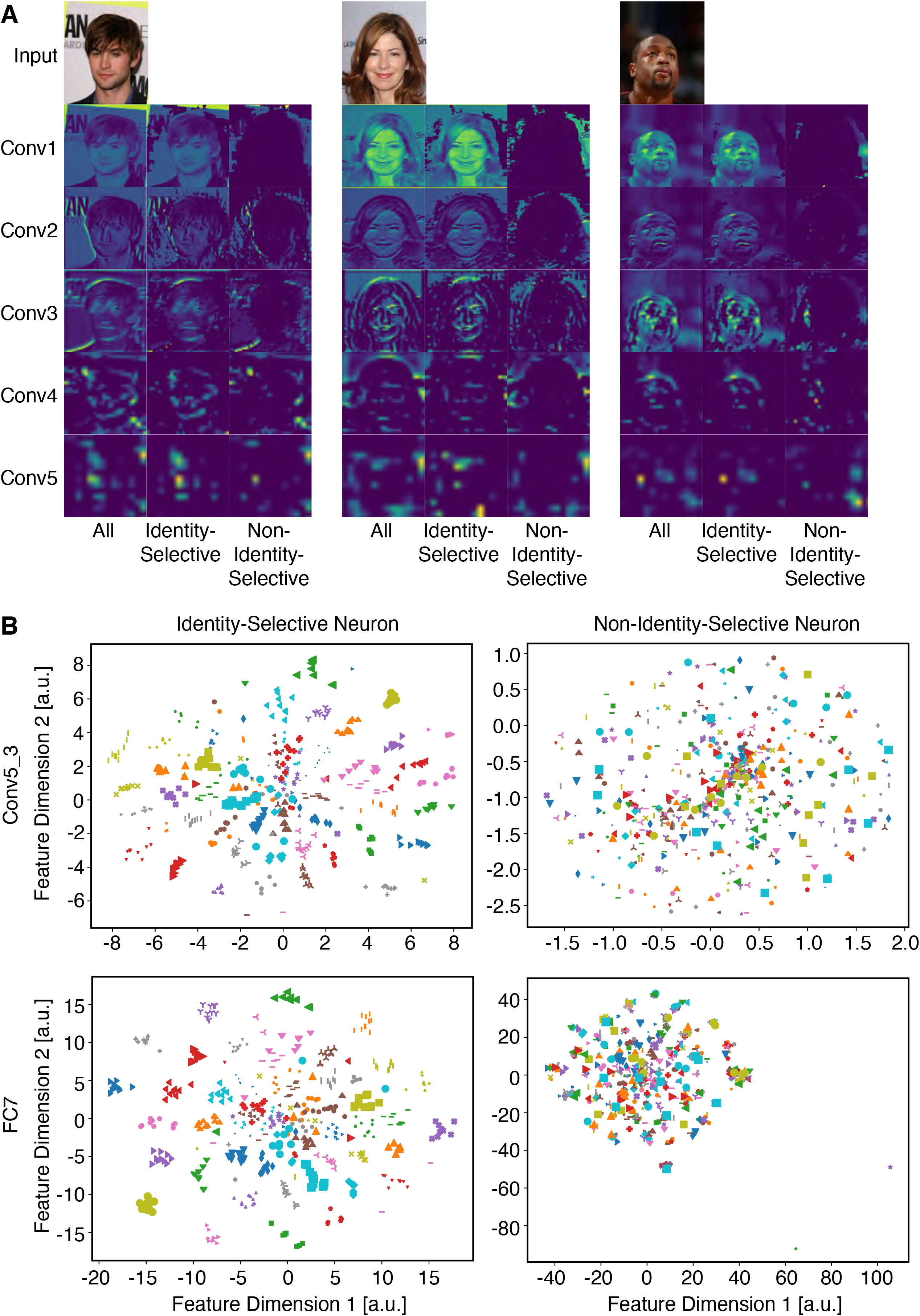
Visualization of the deep neural network (DNN). **(A)** Example activation maps for a subset of DNN layers. Left: all neurons. Middle: identity-selective neurons. Right: non-identity-selective neurons. **(B)** Visualization in the t-SNE space for DNN layers Conv5_3 (upper row) and FC7 (lower row). Each color-shape combination represents a face identity (10 faces per identity). The feature dimensions are in arbitrary units (a.u.). Left column: identity-selective neurons. Right column: non-identity-selective neurons.

### Lesion and perturbation of the network

We next investigated how critical the identity-selective neurons as well as the trained network weight structure were to discriminate face identities by lesioning and perturbing the network. First, following each convolutional layer, we added a “RandDrop” layer (i.e., a binary mask applied to the preceding layer; **Fig. 4A**) to randomly set a subset of DNN neurons to be 0, which partially lesioned the network. Indeed, we found that identity recognition accuracy decreased as a function of increasing lesion amount (**Fig. 4B-F**). With a small amount (10%) of information loss (**Fig. 4B**), the network could still well discriminate face identities and had a comparable performance as the intact network (**Fig. 2**). When 30%-50% of DNN neurons were dropped (**Fig. 4C, D**), only earlier layers had a comparable performance as the intact network but later layers had a significant decrease of performance. When 70%-90% of DNN neurons were dropped (**Fig. 4E, F**), the network could not perform identity discrimination any more. Notably, in all these cases, identity-selective neurons still played the dominant role in discriminating the identities and non-identity-selective neurons did not contribute to the discrimination. It is also worth noting that we dropped the same percentage of neurons for every layer, so information loss would accumulate in later layers. Interestingly, we found that dropping neurons from a single layer had only limited impact on recognition performance in subsequent layers (**Fig. S2**; 30% of neurons were dropped; only dropping neurons from the very first layer Conv1_1 impacted performance in subsequent layers), suggesting that the network had great plasticity.

**Fig. 4.**
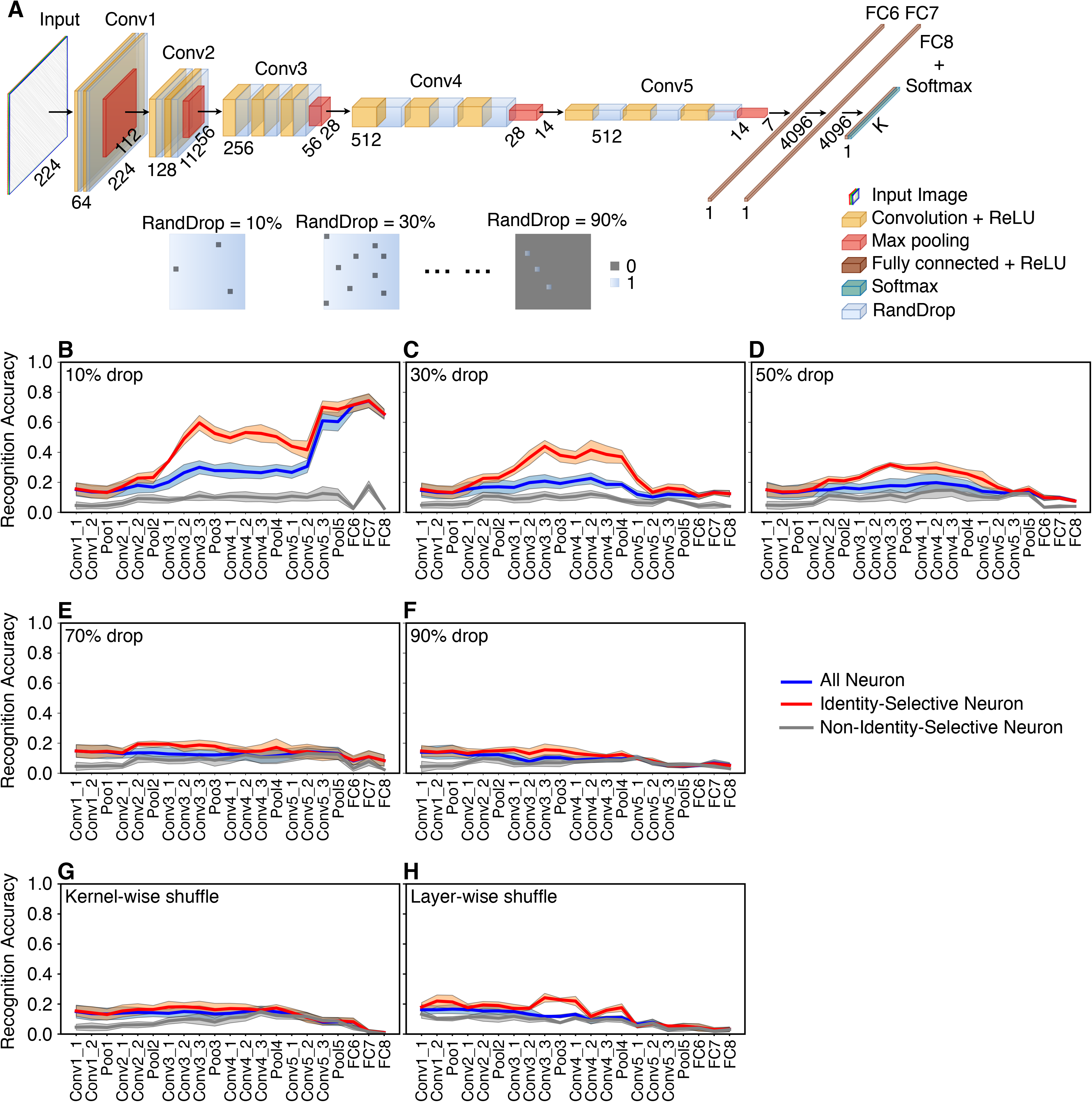
Manipulation of the deep neural network (DNN). **(A)** Illustration of how random dropout of DNN neurons was performed. We added a binary mask following every convolutional layer in the original DNN architecture to randomly deactivate a subset of DNN neurons. The percentage of dropped DNN neurons was controlled by the percentage of zeros in the binary mask and varied from 10% to 90%. **(B-F)** Recognition accuracy following a random dropout of DNN neurons. **(G)** Recognition accuracy following kernel-wise shuffle. **(H)** Recognition accuracy following layer-wise shuffle. Legend conventions as in **Fig. 2**.

Second, we perturbed the network by breaking the optimal weight structure (i.e., connection between DNN neurons) derived from training and generated a random permutation of weights (i.e., connections between DNN neurons) to evaluate the impact of the model training on identity recognition. We employed two approaches. Kernel-wise shuffle (**Fig. 4G**) randomly permuted the weights in a single kernel. Since the kernel size of the network was 3 by 3 for all layers, kernel-wise shuffle permuted the 9 weight values for each kernel. We found that the DNN could not discriminate face identities any more following kernel-wise shuffle (**Fig. 4G**). Unlike kernel-wise shuffle that only rearranged the weights within one kernel, layer-wise shuffle pooled the weights of all kernels from a layer and reorganized the weights to form new kernels. Again, we found that the DNN’s ability to discriminate face identities was abolished (**Fig. 4H**).

Together, our model manipulation suggested that identity-selective neurons as well as the optimal DNN neuron connections derived from training were critical to face identity discrimination.

### Establishing the relationship between artificial DNN neurons and real human neurons

The DNN performs the face recognition task similarly as humans. Does the ensemble of DNN neurons share representational similarity with the ensemble of human neurons? In order to answer this question, we used the same stimuli (500 natural face images of 50 celebrities) and recorded from 490 neurons in the MTL (242 neurons from the amygdala, 186 neurons form the anterior hippocampus, and 62 neurons from the posterior hippocampus; firing rate > 0.15 Hz) of 5 neurosurgical patients (16 sessions in total) [24]. Patients performed a one-back task (**Fig. 5A**; accuracy = 75.7±5.28% [mean±SD across sessions]) and they could well recognize the faces [24]. The responses of 46/490 neurons (9.39%) differed between different face identities in a window 250-1000 ms following stimulus onset and these neurons were the real human identity-selective neurons.

**Fig. 5.**
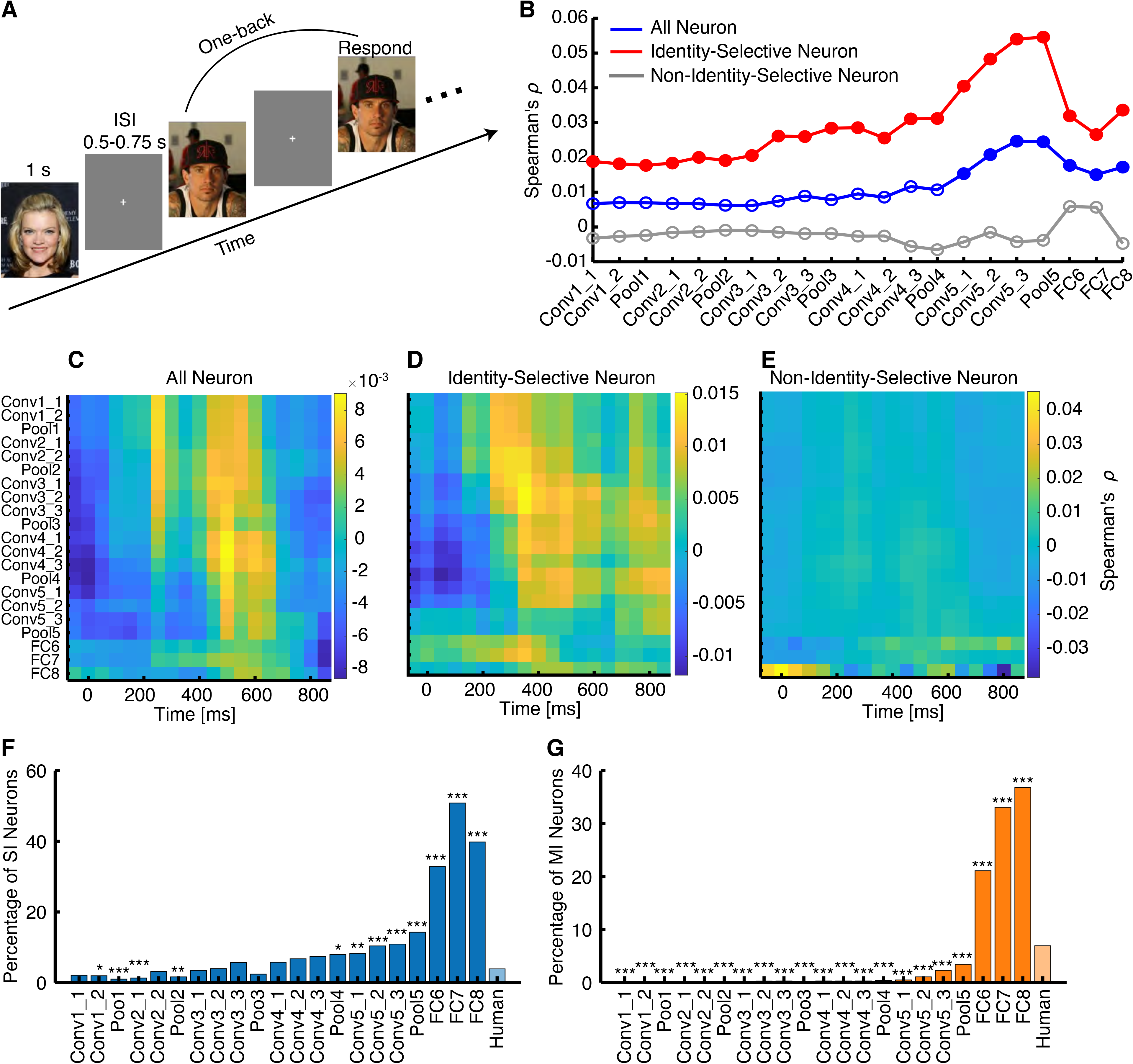
Comparison between the deep neural network (DNN) and real human neurons. **(A)** Task used to acquire single-neuron responses from humans. We employed a one-back task, in which patients responded whenever an identical famous face was repeated. Each face was presented for 1 s, followed by a jittered inter-stimulus-interval (ISI) of 0.5 to 0.75 s. **(B)** Correlation between pairwise distance in the human medial temporal lobe (MTL) neuronal face space and pairwise distance in the DNN face space. Here we used the mean firing rate in a time window 250 ms to 1000 ms after stimulus onset as the response to each face, and we averaged the responses to 10 faces for each face identity. Blue: pairwise distance calculated using all human neurons and all DNN neurons. Red: pairwise distance calculated using human identity-selective neurons (SI and MI neurons) and DNN identity-selective neurons. Gray: pairwise distance calculated using human non-identity-selective neurons and DNN non-identity-selective neurons. Solid circles represent a significant correlation (Spearman’s correlation: P < 0.05, corrected by false discovery rate [FDR] [61] for Q < 0.05) and open circles represent a non-significant correlation. **(C-E)** Temporal dynamics of correlation of pairwise distance between human neurons and DNN neurons (bin size = 500 ms, step size = 50 ms). Color coding indicates the Spearman’s correlation coefficient. **(C)** All neurons. **(D)** Identity-selective neurons. **(E)** Non-identity-selective neurons. **(F)** Percentage of SI neurons in each DNN layer and comparison with neurons from the human MTL. **(G)** Percentage of MI neurons in each DNN layer and comparison with neurons from the human MTL. Asterisks indicate a significant difference in the proportion of neurons using χ^2^-test with Bonferroni correction for multiple comparisons. *: P < 0.05, **: P < 0.01, and ***: P < 0.001.

We first used the mean firing rate in a time window 250-1000 ms after stimulus onset as the response to each face for human neurons (i.e., the same time window as the selection of human identity-selective neurons). We used DNN neurons to construct a face space and correlated that with the human neuronal face space using a distance metric (see **Methods**). Using pairwise activation similarities [21], we found that neuronal pairwise distance of human neurons significantly matched that from the later DNN layers (**Fig. 5B**), suggesting that the population of human neurons encoded the geometry of the face space similarly as DNN neurons. Such match between face spaces was primarily driven by identity-selective neurons: identity-selective neurons showed a significant match across all layers (notably peaked at later DNN layers) whereas non-identity-selective neurons did not show any significant match across layers (**Fig. 5B**). We further investigated the temporal dynamics of the correspondence between face spaces (**Fig. 5C-E**). Again, we found that the match between the DNN and human neuronal face spaces (**Fig. 5C**) was primarily driven by identity-selective neurons (**Fig. 5D**) but not non-identity-selective neurons (**Fig. 5E**). Moreover, identity-selective neurons seemed to match the earlier DNN layers in an earlier time window (**Fig. 5D**).

In addition, we found that compared to the human MTL, the DNN had a significantly higher percentage of SI neurons in later layers (starting from the layer Pool4; **Fig. 5F**; χ^2^-test with Bonferroni correction for multiple comparisons). The DNN also had a significantly higher percentage of MI neurons in the layers FC6, FC7, and FC8 (**Fig. 5G**) but a significantly lower percentage of MI neurons in all other layers, where we did not expect to observe MI neurons because faces of the same identity were not yet clustered.

Together, for the first time, using human single-neuron recordings we directly compared identity selectivity between artificial neurons and real human neurons. Our results have revealed a systematic correspondence between the two face recognition systems.

### Analysis of the region-based feature coding in DNN neurons

We have previously shown a novel region-based feature coding by real neurons in the human MTL [24], i.e., neurons encode identities that are clustered in the feature space (in other words, neurons encode visually similar identities). Here, we explored whether DNN neurons also demonstrated region-based feature coding. We focused on the DNN layers FC6 and FC7, where faces of the same identity was clustered; and we primarily observed human MTL neurons demonstrating region-based feature coding in the face feature spaces from these layers [24]. Notably, the face feature spaces showed an organized structure: for example, Feature Dimension 2 represented a gender dichotomy, and darker skinned faces were clustered at the bottom left corner of the feature space (**Fig. 6A**). Indeed, we found that a large number of MI neurons in these layers (24.9% for FC6 and 36.5% for FC7) encoded identities that were adjacent in the feature space (see **Fig. 6B, C**), demonstrating region-based feature coding. We refer to this subpopulation of MI neurons as feature MI neurons. On the other hand, non-feature MI neurons did not have selective identities clustered in the feature space (**Fig. 6D**), and non-identity-selective neurons did not encode any particular identities (**Fig. 6E**).

**Fig. 6.**
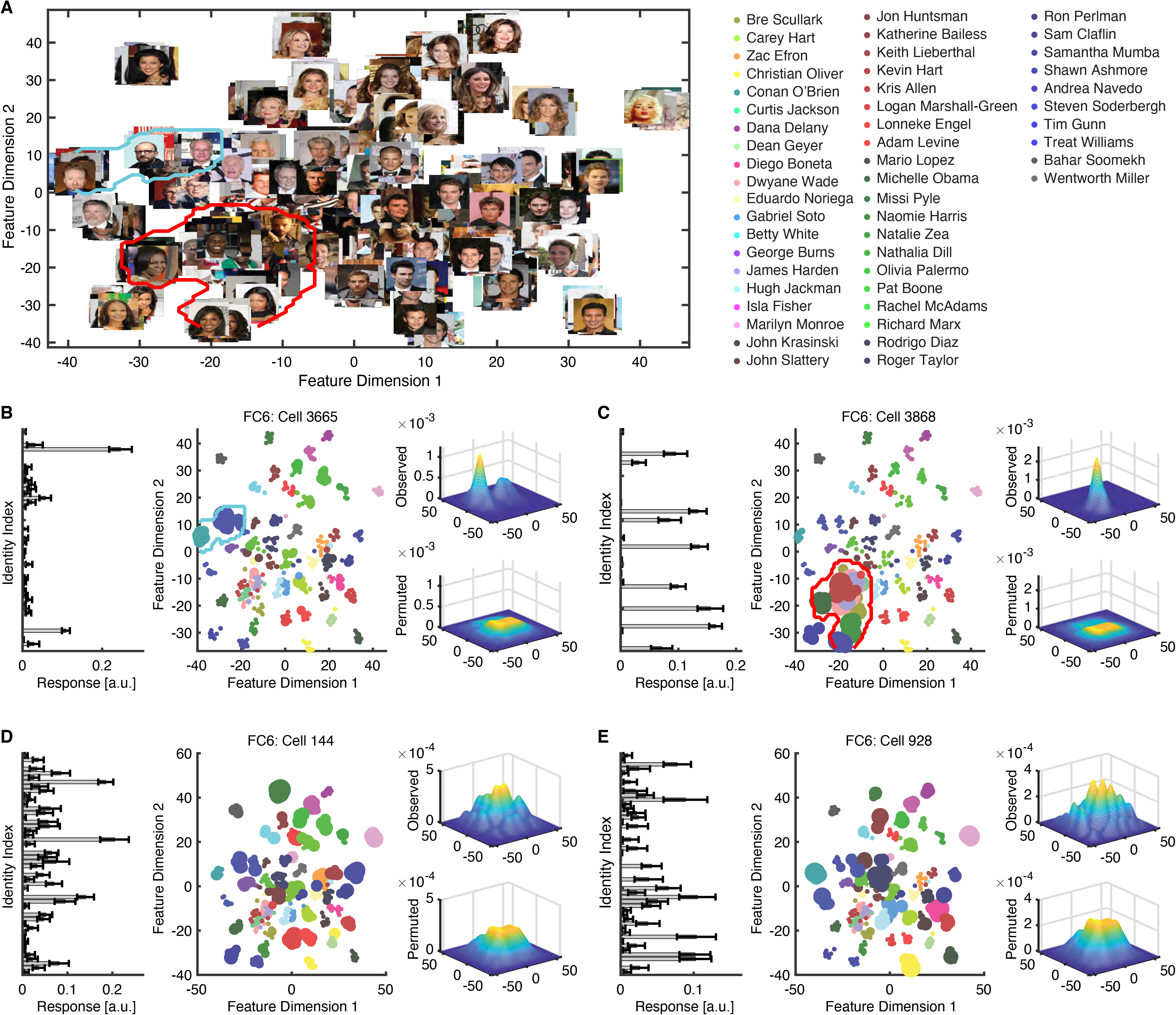
Region-based feature coding in DNN neurons. **(A)** The face feature space constructed by t-distributed stochastic neighbor embedding (t-SNE) for the DNN layer FC6. All stimuli are shown in this space. **(B-E)** Example DNN neurons. **(B, C)** Two example MI neurons that demonstrated region-based feature coding (i.e., the neurons encoded a region in the feature space). **(D)** An example MI neuron that did not demonstrate region-based feature coding (i.e., the encoded identities were not adjacent to each other in the feature space). **(E)** A non-identity-selective neuron (i.e., the neuron did not encode any particular identities). All examples were from the DNN layer FC6. (Left) DNN neuron responses to 500 faces (50 identities; grouped by individual identity) in arbitrary units (a.u.). Error bars denote ±SEM across faces. (Middle) Projection of the DNN activation onto the feature space. Each color represents a different identity (names shown in the legend). The size of the dot indicates the level of activation. (Right) Estimate of the spike density in the feature space. By comparing observed (upper) vs. permuted (lower) responses, we could identify a region where the observed neuronal response was significantly higher in the feature space. This region was defined as the tuning region of a neuron (delineated by the red/cyan outlines).

At the population level, we found that the tuning region of an individual feature MI neuron covered approximately 5-6% of the 2D feature space (**Fig. 7A**; note that when we calculated the tuning region, we adjusted the kernel size to be proportional to the feature dimensions such that the percentage of space coverage was not subject to the actual size of the feature space). In contrast, the response of an individual SI or non-feature MI neuron covered a significantly smaller region in the feature space (**Fig. 7A**; two-tailed unpaired *t*-test: P < 0.001 for all comparisons). As expected, the distance in the face space between encoded identities was smaller for feature MI neurons compared with non-feature MI neurons (**Fig. 7B**). As a whole, the entire DNN neuronal population covered approximately 55-60% of the feature space (**Fig. 7C**; some areas were encoded by multiple neurons), and the covered areas were similar for SI, feature MI, and non-feature MI neurons (**Fig. 7C**).

**Fig. 7.**
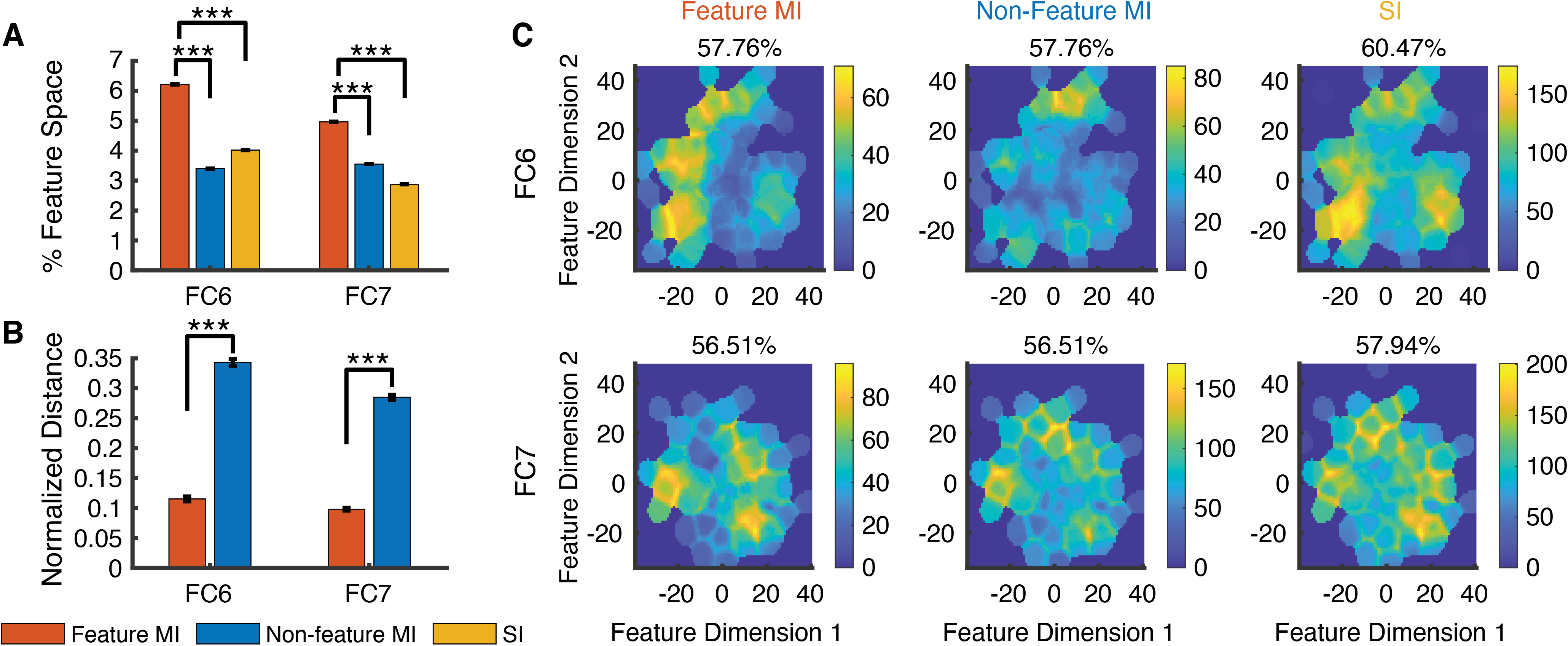
Summary of region-based feature coding for SI and MI neurons. **(A)** Percentage of feature space covered by tuning regions of SI and MI neurons. Note that here we did not apply the threshold for minimal cluster size for SI and non-feature MI neurons in order to compare between different types of neurons. **(B)** Normalized distance between MI neuron’s selective identities in the feature space. To be comparable for different layers, Euclidean distance was normalized by the maximum distance (i.e., diagonal line) of the feature space. Error bars denote ±SEM across neurons. Asterisks indicate a significant difference between feature MI neurons and non-feature MI neurons using two-tailed unpaired *t*-test. ***: P < 0.001. **(C)** The aggregated tuning regions of the neuronal population. Color bars show the counts of overlap between individual tuning regions. Numbers above the density map show the percentage of feature space covered by the tuning regions of the neuronal population.

Together, we found that as neurons in the human MTL, DNN neurons also demonstrated region-based feature coding. Region-based feature coding may provide a possible mechanism that explains how the DNN performs face recognition.

## Discussion

In this study, we analyzed the response characteristics of a face recognition DNN and found that identity-selective neurons in the DNN could generalize their discriminability to face identities shown in various styles as well as face identities that were not involved in the training. By establishing the coding similarity with real human neurons, our study provided an important method to understand face coding in humans. Furthermore, by analyzing an artificial neural network dedicated to face recognition, we will be able to formulate hypotheses that can be validated in future studies.

### Possible caveats

Although the SVM had substantially more features (i.e., DNN neurons used for classification) than observations (i.e., training faces), the high recognition accuracy of identity-selective neurons in testing suggested that our results could not be simply explained by overfitting. Furthermore, the low recognition accuracy of non-identity-selective neurons in testing provided specificity of our approach; and notably, later DNN layers had fewer neurons but a higher recognition accuracy. Interestingly, such “overfitting” also appears in the human brain as a large number of neurons are often simultaneously activated by a single percept; and the theory of backward feature correction can well explain such “overfitting” [34].

We identified the DNN neurons critical for identity recognition, and the evolution of identity-selective neurons across DNN layers (**Fig. 1B**). The identity-selective neurons in the earlier layers primarily corresponded to image pixels containing information about faces, but with the increase of kernel size in later layers, identity-selective neurons encoded more holistic information about face identities. In particular, the fully connected layers utilized information from all neurons from the previous layers. Therefore, identity selectivity could not be solely attributed to the receptive field of the DNN neurons.

In this study, we found that although cartoon faces had a decreased discriminability in general, they had a similar pattern of response as natural faces across DNN layers (**Fig. 2A, C-F**). Furthermore, we found that inverted faces elicited a similar response as upright faces in the early and middle DNN layers (**Fig. 2A, I**), suggesting that the DNN used similar information for both inverted and upright faces, a form of viewpoint invariance. However, in contrast to all upright faces that had increasing discriminability across layers, the discriminability decreased in later layers for inverted faces. Therefore, the impaired discriminability of inverted faces in humans may stem from neurons downstream in the visual processing stream [33]. Furthermore, we found that the response of face identity-selective neurons could well generalize within faces but barely generalize to non-face objects, consistent with a dedicated and specialized face perception system [1, 23]. Lastly, consistent with our DNN lesion results (**Fig. S2**), it has been shown that DNN neurons demonstrate distributed and sparse codes to represent different face attributes [26].

### Comparison between artificial and human neurons

We found that identity-selective neurons had a general discriminability to face identities shown in various styles, consistent with feature-invariant coding of face identities by neurons in the human medial temporal lobe (MTL) [6, 7]. Furthermore, identity selectivity could be generalized to face identities that were not involved in the training, similar to how memory is formed in the human brain [35]. Consistent with our prior findings that only a small proportion (~20%) of human neurons are involved in coding a certain task aspect, such as emotion content [36], emotion subjective judgment [37], attention [38, 39], task sequence [39], visual selectivity [38], eye movement [40], social judgment [41], as well as face identity [24], in the present study we found a large population non-identity-selective DNN neurons that did not contribute to coding face identities. We further quantitatively compared the proportion of SI and MI neurons between the DNN and human brain. Notably, the distribution of identity-selective and non-identity-selective DNN neurons across layers may provide new insights into understanding the human visual processing stream, where our currently available technology does not allow simultaneous sampling of neurons along the entire visual processing stream.

Notably, we found that DNN neurons demonstrated region-based feature coding as neurons in the human MTL [24], a novel and important mechanism that may explain how the DNN performs face recognition. It has been shown that the higher visual cortex encodes visual features of the stimuli [10–13], analogous to feature extraction and representation in the DNN, while the MTL encodes semantic categories independent of visual features [6, 7], analogous to the output of the DNN. Region-based feature coding, which shows that DNN neurons in the later layers encode a region in the feature space using information from the earlier layers and are selective to identities that fall in this region, therefore bridges the representation of image features in the earlier layers and the representation of semantic categories (e.g., face identities) in the later layers. Such processing hierarchy is similar to how the brain transitions from a perception-driven representation of features in the higher visual cortex to a memory-driven representation of semantics in the MTL. Region-based feature coding may also provide an account for object recognition (e.g., using the AlexNet) and visual selectivity (also observed in human MTL neurons): objects falling within the coding region of a neuron may elicit an elevated response. Again, this mechanism applies to both artificial and human neurons.

### Contribution of DNNs to understanding visual processing

It is worth noting that most previous studies of face space had to use computer-generated faces in order to parametrically vary the faces [10, 42, 43]. Rapid advances in computer vision and development of DNNs have provided an unprecedented opportunity to extract features from *real* human faces and subsequently manipulate these features to generate *new* unique faces while providing well controlled stimuli to investigate differences in neural responses to feature changes. Inspired by the primate visual system, DNNs have made impressive progress on the complex problem of recognizing faces across variations of camera distance, viewpoint, illumination, and changes in expression. It has been widely used to study face recognition and representation (see [44] for a review). DNNs may help researchers to understand the functional architecture of human brains and incorporating neuroscience findings into DNNs can test the computational benefits of fundamental organizational features for the visual system [45]. The DNNs do share similarities with the primate visual system and can therefore help us to better understand visual processing in primates [46]. For example, within a class of biologically plausible hierarchical neural network models, there is a strong correlation between a model’s categorization performance and its ability to predict single-neuron responses from the IT cortex [47]. In addition, both human neuroimaging studies [17] and intracranial electroencephalogram (EEG) studies [48] have shown that features in DNNs can be represented in the human brain, which in turn can explain our ability to recognize individual faces. Notably, recent studies in monkeys have shown that images synthesized by DNN can control neural population activity [11] and evolution of synthetic images given online neural feedback can even drive neural response to the maximum [12].

### Future directions

It is worth noting that most faces in the present study had a frontal view and were primarily emotionally neutral. A future study will further test faces from different angles (e.g., profile faces) and/or emotional faces. Furthermore, an interesting future study will be to compare DNN lesion results with behavior of human brain lesion patients, who demonstrate impaired face perception (e.g., [36]).

A central challenge in cognitive neuroscience is to understand how the brain encodes faces. In particular, it remains largely unclear how visual experience, learning, and memory shape face perception and recognition. There are two competing hypotheses about the emergence of face-selective neurons. One hypothesis argues that face-selective neurons require visual experience to develop, and this hypothesis has been supported by fMRI studies in the monkey fusiform face areas [49]. The other hypothesis argues that face-selective neurons have an innate origin, and this hypothesis has been supported by studies from human infants and adults without visual experience of faces [50–53]. Along this line, a future study will be needed to explore if identity-selective neurons can spontaneously emerge from untrained neural network. Although the DNN used in the present study was a pre-trained ANN, it demonstrated strong ability to generalize to new faces, comparable to the human visual system. Importantly, *in silico* experiments allow us to test a large set of parameters and better control experimental conditions, which is often not feasible when directly working with human or animal subjects. Together, our present study not only highlights the importance of the direction of training and visual experience in shaping the neural response to face identities, but also provides a useful approach to test these hypotheses and reconcile previous findings.

## Methods

### Stimuli

We employed the following stimuli in this study (**Fig. 2**).

1. For the original stimuli, we used faces of celebrities from the CelebA dataset [54], and we selected 50 identities with 10 images for each identity, totaling 500 face images. The identities were selected to include both genders and multiple races (see also **Fig. 6A**).
2. We selected another 500 faces from 50 different identities (10 images per identity) from the CelebA dataset as a testing set.
3. We generated four versions of cartoon faces (Hayao, Hosoda, Paprika, Shinkai) of the original stimuli using CartoonGAN [55].
4. We generated Mooney faces by first transforming the original images into gray scale. We then filtered the images with a two-dimensional Gaussian smoothing kernel with standard deviation of 0.5. We lastly thresholded the images using a threshold determined for each individual face based on its luminance (threshold = mean luminance of the cropped image center − 0.03).
5. We randomly selected 500 caricature faces of 50 identities (10 images per identity) from the IIIT-CFW dataset [56].
6. We randomly selected 500 non-faces objects from 50 categories (10 objects per category) from the ImageNet database [57].

### Deep neural network (DNN)

We used the well-known deep neural network (DNN) implementation based on the VGG-16 convolutional neural network (CNN) architecture [15] (see **Fig. 1A** for details). We fine-tuned the layer FC8 with the original CelebA stimuli to confirm that this pre-trained model was able to discriminate the identities (accuracy > 90%). To visualize the DNN response, we subsequently applied a t-distributed stochastic neighbor embedding (t-SNE) method to convert high-dimensional features into a two-dimensional feature space. t-SNE is a variation of stochastic neighbor embedding (SNE) [58], a commonly used method for multiple class high-dimensional data visualization [59]. We applied t-SNE for each layer, with the cost function parameter (Prep) of t-SNE, representing the perplexity of the conditional probability distribution induced by a Gaussian kernel, set individually for each layer. We also used t-SNE to construct a face feature space so that we were able to investigate region-based feature coding for both DNN and human neurons [24].

### Selection of identity-selective neurons

To select identity-selective neurons, we used a one-way ANOVA to identify identity-selective neurons that had a significantly unequal response to different identities (P < 0.01; **Fig. S1A**). We further imposed an additional criterion to identify a subset of identity-selective neurons with selective identities (**Fig. S1A**): the response of an identity was 2 standard deviations (SD) above the mean of responses from all identities. These identified identities whose response stood out from the global mean were the encoded identities. We refer to the neurons that encoded a single identity as single-identity (SI) neurons and we refer to the neurons that encoded multiple identities as multiple-identity (MI) neurons.

### Single-neuron recordings in human neurosurgical patients

To acquire the neuronal response from humans, we conducted single-neuron recordings from 5 neurosurgical patients (16 sessions in total). All participants provided written informed consent using procedures approved by the Institutional Review Board of West Virginia University (WVU). The detailed procedure has been described in our previous study [24]. Briefly, we employed a 1-back task for the original CelebA stimuli (**Fig. 5A**). In each trial, a single face was presented at the center of the screen for a fixed duration of 1 second, with uniformly jittered inter trial interval (ITI) of 0.5-0.75 seconds. Patients pressed a button if the present face image was *identical* to the immediately previous image. Each face was shown once unless repeated in one-back trials; and we excluded responses from one-back trials to have an equal number of responses for each face.

We recorded from implanted depth electrodes in the amygdala and hippocampus from patients with pharmacologically intractable epilepsy. Bipolar wide-band recordings (0.1-9000 Hz), using one of the eight microwires as reference, were sampled at 32 kHz and stored continuously for off-line analysis with a Neuralynx system. The raw signal was filtered with a zero-phase lag 300-3000 Hz bandpass filter and spikes were sorted using a semi-automatic template matching algorithm as described previously [60]. Units were carefully isolated and recording and spike sorting quality were assessed quantitatively. Only units with an average firing rate of at least 0.15 Hz (entire task) were considered. Only single units were considered. Trials were aligned to stimulus onset and we used the mean firing rate in a time window 250 ms to 1000 ms after stimulus onset as the response to each face.

### Pairwise distances in the face space

We further employed a pairwise distance metric [21] to compare neural coding of face identities between human and DNN neurons. For each pair of identities, we computed two distance metrics. The human neuronal distance metric was the Euclidean distance between firing rates of all recorded neurons. The DNN distance metric was the Euclidean distance between feature weights of all DNN neurons. We then correlated the human neuronal distance metric and the DNN distance metric. A significant correlation indicated that the human neuronal face space matched the DNN face space [21]. We computed the correlation for each DNN layer so that we could determine the specific layer that the neuronal population encoded. For each face identity, we averaged the response of all faces of that identity to get one mean firing rate. To get temporal dynamics, we used a moving window with a bin size of 500 ms and a step size of 50 ms. The first bin started −300 ms relative to trial onset (bin center was thus 50 ms before trial onset), and we tested 19 consecutive bins (the last bin was thus from 600 ms to 1100 ms after trial onset).

### Selection of feature neurons

We employed the same procedure to select DNN feature neurons as we did with human neurons [24]. We first estimated a continuous spike density map in the feature space by smoothing the discrete activation map using a 2D Gaussian kernel (kernel size = feature dimension range * 0.2, SD = 4). We then estimated statistical significance for each pixel by permutation testing: in each of the 1000 runs, we randomly shuffled the labels of faces. We calculated the p-value for each pixel by comparing the observed spike density value to those from the null distribution derived from permutation. We lastly selected the region with significant pixels (permutation P < 0.01, cluster size > 0.23 * pixel number in the whole space). We also applied a mask to exclude pixels from the edges and corners of the spike density map where there were no faces because these regions were susceptible to false positives given our procedure. If a neuron had a region with significant pixels, the neuron was defined as a “feature neuron” and demonstrated “region-based feature coding”. We selected feature neurons for each individual DNN layer.

## Data availability

All data and statistical analysis code are available on GitHub (https://github.com/JingeW/ID_selective).

## Acknowledgements

We thank all patients for their participation and staff from WVU Ruby Memorial Hospital for support with patient testing. This research was supported by an NSF CAREER Award (1945230), Air Force Young Investigator Program Award (20RT0829), Dana Foundation Clinical Neuroscience Award, ORAU Ralph E. Powe Junior Faculty Enhancement Award, West Virginia University (WVU), and WVU PSCoR Program (to S.W.), and an NSF Grant (OAC-1839909) and the WV Higher Education Policy Commission Grant (HEPC.dsr.18.5) (to X.L.). The funders had no role in the study design, data collection and analysis, decision to publish, or preparation of the manuscript.

## Author Contributions

J.W., R.C., X.L., and S.W. designed research. J.W., R.C., and S.W. performed experiments. N.J.B. performed surgery. J.W., R.C., and S.W. analyzed data. J.W., X.L., and S.W. wrote the paper. All authors discussed the results and contributed towards the manuscript.

## Supplemental Figure Legends

**Fig. S1.**
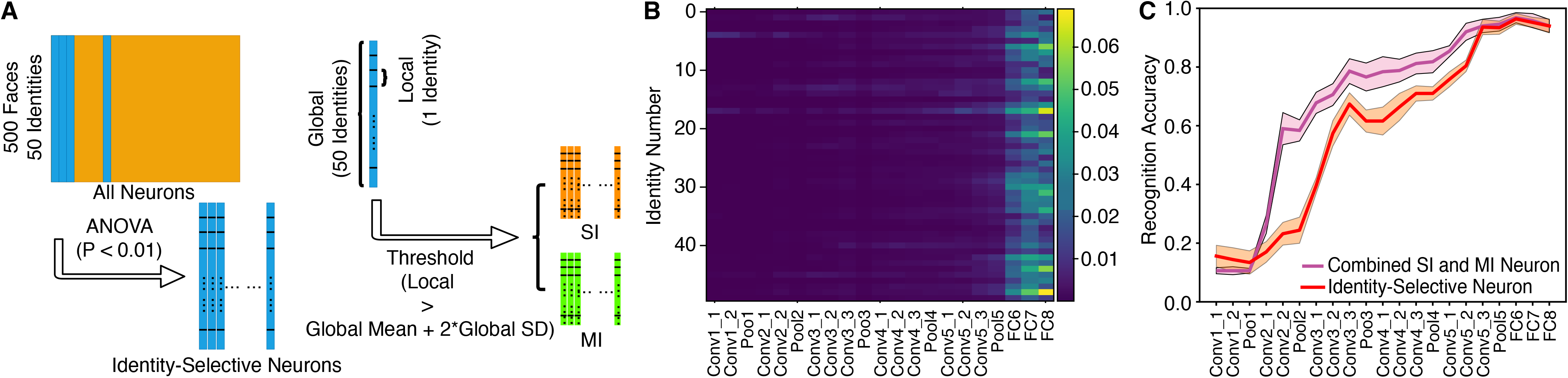
Additional results for identity-selective neurons. **(A)** Procedure for selecting identity-selective neurons. We used a one-way ANOVA to identify identity-selective neurons that had a significantly unequal response to different identities (P < 0.01). We further imposed an additional criterion to identify a subset of identity-selective neurons with selective identities: the response of an identity was 2 standard deviations (SD) above the mean of responses from all identities. These identified identities whose response stood out from the global mean were the encoded identities. We refer to the neurons that encoded a single identity as single-identity (SI) neurons and we refer to the neurons that encoded multiple identities as multiple-identity (MI) neurons. **(B)** The encoding frequency of each face identity in each deep neural network (DNN) layer. Color coding indicates the encoding frequency of each DNN layer (i.e., the number of DNN neurons encoding an identity divided by the total number of DNN neurons of a layer). **(C)** Combined SI and MI neurons demonstrated even better discriminability of face identities than identity-selective neurons. Identity recognition accuracy is shown for each DNN layer. Error shade denotes one standard deviation across 5-fold cross validation. Magenta: combined SI and MI neurons. Red: identity-selective neurons.

**Fig. S2.**
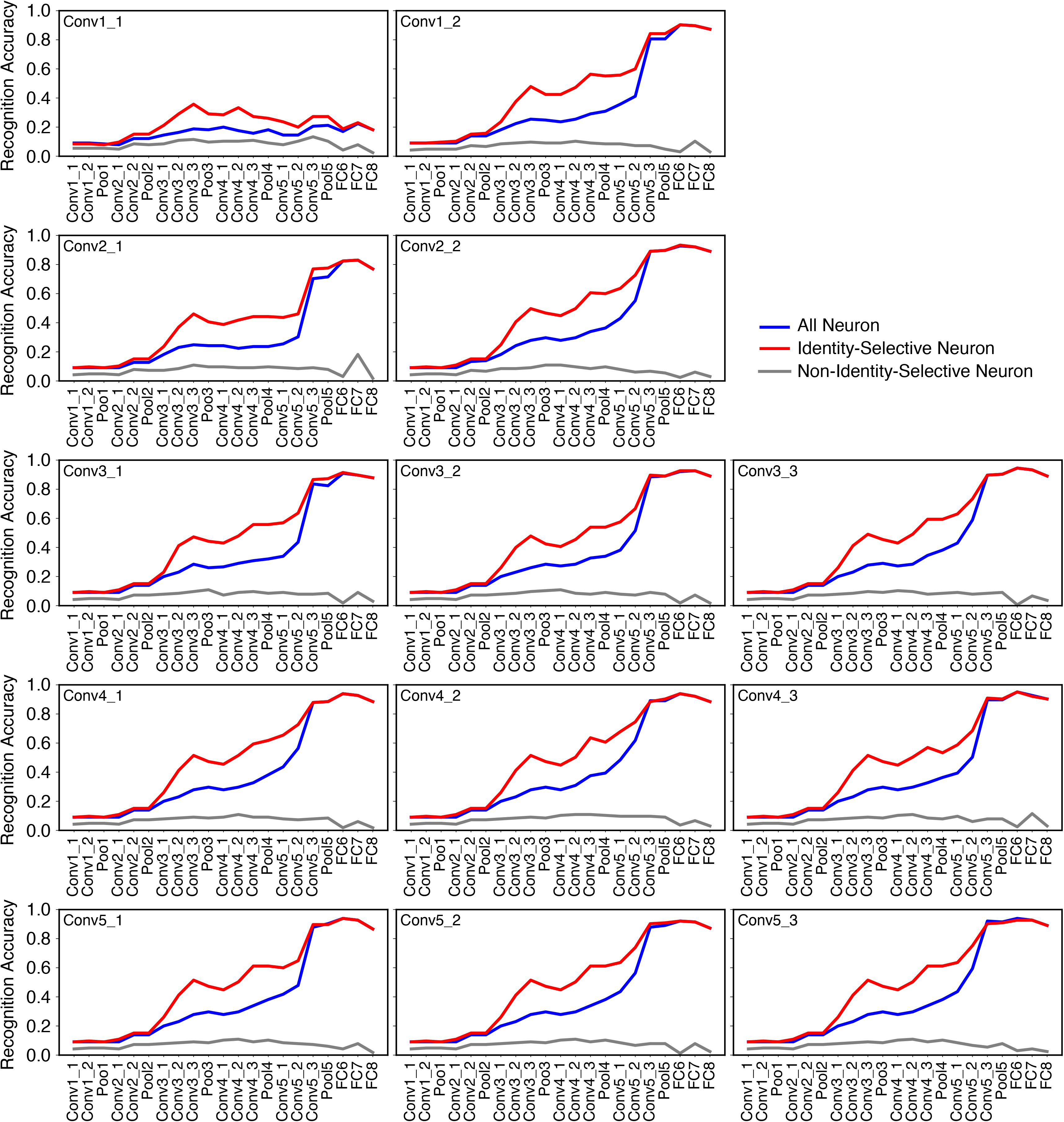
Recognition accuracy following a random dropout of a single layer of DNN neurons. 30% of neurons were dropped. The label in each plot indicates the layer that dropout was performed. Legend conventions as in **Fig. 2**.

